# *Ex Vivo* Validation of Six FDA-Approved Non-Receptor Tyrosine Kinase Inhibitors (NRTKIs) as Antivirals to Pandemic and Seasonal Influenza A Viruses

**DOI:** 10.1101/2022.01.19.476993

**Authors:** Robert Meineke, Sonja Stelz, Maximilian Busch, Christopher Werlein, Mark Kühnel, Danny Jonigk, Guus F. Rimmelzwaan, Husni Elbahesh

**Affiliations:** Research Center for Emerging Infections and Zoonoses (RIZ), University of Veterinary Medicine (TiHo), Bünteweg 17, 30559 Hannover, Germany; Department of Pathology, Hannover Medical School (MHH), Carl-Neuberg-Straße 1, 30625 Hannover, Germany; Member of the German Center for Lung Research (DZL), Biomedical Research in Endstage and Obstructive Lung Disease Hannover (BREATH)

## Abstract

Influenza viruses are important respiratory pathogens that cause substantial morbidity and mortality annually. In addition to seasonal influenza outbreaks, new emerging influenza A viruses (IAV) can cause pandemic influenza outbreaks. Apart from effective vaccines, there is a need for better treatment options to combat infections with these viruses when vaccines are not available or show reduced efficacy (e.g., in immmunocompromised patients). The limited range of licensed antiviral drugs and emergence of drug-resistance mutations highlight the need for novel intervention strategies like host-targeted antivirals. Repurposing FDA-approved kinase inhibitors may offer a fast-track for a new generation of host-targeted antivirals. Small molecule kinase inhibitors (SMKIs) can inhibit replication of viruses and improve survival *in vivo*; however, no SMKI has been approved for clinical use against IAV infections. In the present study, we tested eight non-receptor tyrosine kinase-inhibitors (NRTKIs) used to treat cancer and autoimmune diseases for their antiviral potential. Six of those potently inhibited virus replication (*≥*1,000-fold) in A549 cells infected with either A(H1N1)pdm09 or seasonal A(H3N2) strains. These compounds were validated in a biologically relevant *ex vivo* model of human precision-cut lung slices (hPCLS) to provide proof of principle and show efficacy against contemporary seasonal and pandemic IAVs. We identified the steps of the virus infection cycle affected by these inhibitors and assessed the effect of these NRTKIs on the host response. Considering their established safety profiles, our studies show that the use of these NRTKI shows promise and warrants further development as an alternative strategy to treat influenza virus infections.

## INTRODUCTION

Influenza viruses are an important cause of respiratory tract infections in humans and are responsible for substantial annual morbidity and mortality, especially in individuals at high risk, like older adults or immunocompromised patients. The most important preventive measure to protect humans from influenza virus infections is vaccination. Currently used vaccines mainly aim at the induction of virus neutralizing antibodies to the viral hemagglutinin (HA). For vaccines to be effective they need to antigenically match the circulating strains. ^1–3^. Due to a lack of proof-reading activity of their RNA-dependent RNA polymerase (RdRp), influenza viruses can accumulate mutations that lead to antigenic drift and allow them to evade recognition by virus-neutralizing (VN) antibodies induced by previous infections or vaccinations. These antigenic changes also necessitate annual update of influenza vaccines against seasonal influenza ^4^. Moreover, emergence of novel influenza viruses that arise through genetic reassortment after interspecies transmission. Emergence of these viruses in human populations that largely lack VN antibodies can result in pandemic outbreaks. At the early stages of a pandemic, effective vaccines are not readily available, as was the case during the pandemic of 2009 caused by swine-origin A(H1N1) influenza A viruses (IAVs).

In the absence of efficacious vaccines, virus-targeted antivirals can offer some protection if administered within the therapeutic window. Until recently, influenza antivirals were comprised of two classes, Adamantanes that target the viral M2 ion channel protein and neuraminidase inhibitors (NAI). Due to almost ubiquitous resistance found in currently circulating strains, Adamantanes have been rendered ineffective and are no longer used in clinical practice ^5^. The levels of currently circulating viruses that carry resistance mutations to NAIs (such as oseltamivir) isolated from otherwise healthy individuals is only ∼4-5% ^6^. A more recent strategy has been to target the influenza polymerase proteins, with Favipiravir (T705), Baloxavir and Pimodivir targeting the viral PB1, PA and PB2 polymerase subunit proteins, respectively being the most widely used antivirals. All seem to inhibit viruses that are resistant to Adamantanes and NAIs ^7, 8^. In 2018, Baloxavir was approved for the treatment of acute and “uncomplicated” influenza infections in the United States^9, 10^. Surprisingly, Uehra et al. found that even in otherwise healthy adults and adolescents the emergence of strains carrying Baloxavir resistance mutations was as high as ∼10% and could potentially be higher in immunocompromised patients ^11, 12^.

The sustained circulation of virus variants resistant to current antivirals and the low genetic barrier to achieve resistance highlight the need for host-targeted therapeutics; that do not suffer from these limitations. Given that all viruses rely on host-cellular machinery for replication at every step of their life cycle, several host proteins have been shown to be required for efficient viral replication and pathogenesis ^13–17^. Host kinases link a myriad of signaling pathways used by viruses; and as such, they offer attractive targets for potential host-directed therapeutics against infections by several viruses including influenza viruses ^13, 14, 18^. Host kinases catalyse phosphorylation of lipids or proteins, at either tyrosine or serine/threonine residues that not only serves to relay and amplify cell signaling, but phosphorylation can also lead to conformational changes in proteins that facilitate protein-protein interactions ^19, 20^.

Although the human kinome consists of more than 550 kinases, currently available kinase inhibitors only target ∼10% of kinases and are primarily used to treat cancers ^21–25^. To date, only 62 small-molecule kinase inhibitors (SMKIs) have been FDA-approved and the majority of these target tyrosine kinases ^26^. Non-receptor tyrosine kinases (NRTKs) are cytoplasmic or membrane-anchored kinases that closely associate with cellular receptors or receptor complexes to mediate outside-in signaling ^26^. NRTKs like most kinases contain a catalytic kinase domain, as well as several protein-protein interaction motifs (e.g., SH2, SH3, PH domains, etc.) necessary to relay cell signals. NRTKs include Abl, FAK, JAK, Src and BTK, all of which have been reported to play a role in IAV infections. Previous studies have demonstrated that NRTK inhibitors (NRTKIs) can modulate pro- and anti-viral signaling *in vitro* and result in reduced viral pathogenesis and increased survival *in vivo* ^13, 16, 27–32^. However, no NRTKIs or SMKIs have been approved for clinical use against influenza. In the present study, we characterized the effect of eight currently available and FDA-approved NRTKIs on IAV replication, six of which were validated using a biologically relevant *ex vivo* model of human precision-cut lung slices (hPCLS). hPCLS maintain near-native structural integrity of lung tissues that allow complex and 3D cell-cell interactions of epithelial cells, mesenchymal tissue as well as the vascular compartment; thereby, offering an advantage over cultured epithelial monolayers ^33–35^. We also identify which steps of the virus infection cycle are affected by specific NRTKIs; thereby providing insights into potential mechanisms of action in the context of influenza virus infections.

## RESULTS

### NTRKI treatment inhibits IAV replication *in vitro*

To identify non-toxic concentrations of our inhibitors, we used the CellTiter-Glo (CTG) assay in which cell-viability is based on ATP content in healthy cells. We identified concentrations that resulted in cell viability ≥90% relative to mock-treated cells (cells treated with 0.1% DMSO). The highest concentration with ≥90% relative viability was defined as the 1x concentration ([1x]_max_) (**Fig. 1/Table 1**).

**Figure 1.**
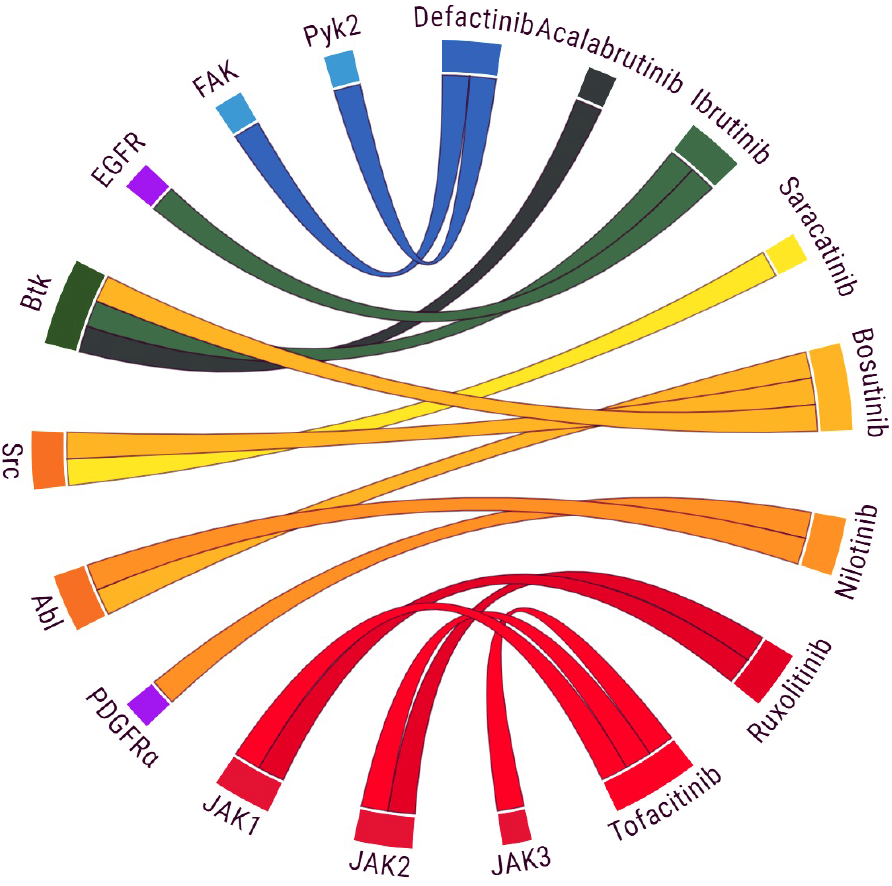
Main target specificity of candidate NRTKIs used in this study.

**Table 1.**
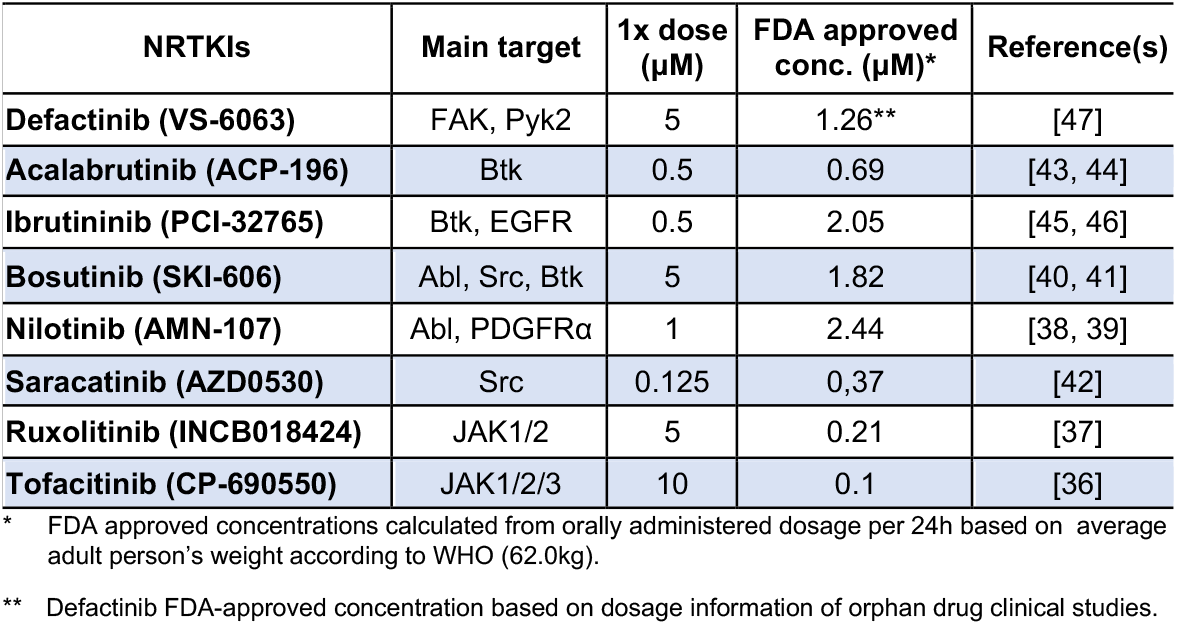
. Main targets of FDA-approved NRTKIs used.

| NRTKIs | Main target | 1x dose (μM) | FDA approved conc. (μM)* | Reference(s) |
| --- | --- | --- | --- | --- |
| Defactinib (VS-6063) | FAK, Pyk2 | 5 | 1.26** | [47] |
| Acalabrutinib (ACP-196) | Btk | 0.5 | 0.69 | [43, 44] |
| Ibrutinib (PCI-32765) | Btk, EGFR | 0.5 | 2.05 | [45, 46] |
| Bosutinib (SKI-606) | Abl, Src, Btk | 5 | 1.82 | [40, 41] |
| Nilotinib (AMN-107) | Abl, PDGFRα | 1 | 2.44 | [38, 39] |
| Saracatinib (AZD0530) | Src | 0.125 | 0.37 | [42] |
| Ruxolitinib (INCB018424) | JAK1/2 | 5 | 0.21 | [37] |
| Tofacitinib (CP-690550) | JAK1/2/3 | 10 | 0.1 | [36] |

\* FDA approved concentrations calculated from orally administered dosage per 24h based on average adult person's weight according to WHO (62.0kg).
\*\* Defactinib FDA-approved concentration based on dosage information of orphan drug clinical studies.

Next, we determined the effect of eight NRTKIs on influenza A virus replication. A549 cells were infected with either pandemic A(H1N1)pdm09 strain A/Netherlands/602/09 (NL09) or seasonal A(H3N2) strain A/Netherlands/241/11 (NL11) at a multiplicity of infection (MOI) of 1 in the presence or absence of [1x, 0.5x and 0.25x]_max_ concentrations of the respective inhibitors following inoculation. Culture supernatants were collected at 2, 24, 48, and 72 hours post infection (hpi) and virus titers were determined by median tissue culture infectious dose (TCID_50_) assay. We observed a dose-dependent reduction of viral titers by six of the eight inhibitors with at least one of the concentrations ranging from 2- to 1,000-fold (**Fig. 2A**) reduction of virus titers. As visualized by the heatmap (**Fig. 2B),** there was variability in both magnitude and duration of the reduction of virus titers, and in general, the effect was more pronounced in NL11 (H3N2) infected cells than NL09 (pH1N1). Although we observed only a transient reduction at 24 hpi with Tofacitinib (TF) (JAK1/2/3 inhibitor)^36^, we did not observe any significant reduction with Ruxolitinib (RX) (JAK1/2 inhibitor)^37^. The highest and most sustained level of reduction was observed with Nilotinib (NI) (Abl/PDGFRa inhibitor)^38, 39^: >1,000-fold (3-log_10_) reduction. Bosutinib (BO) (Abl/Src/Btk inhibitor)^40, 41^ and Saracatinib (SA) (Src inhibitor)^42^ also showed marked inhibition (SA ∼ 5- to 25-fold; BO ∼10- to 1,000-fold). Acalabrutinib (AC) (Btk inhibitor)^43, 44^ had very little effect on NL09 replication but showed a greater and more sustained inhibition (5- to 25-fold) of NL11 replication at the higher concentrations (0.25 and 0.5 μM). Similarly, Ibrutinib (IB) (Btk/EGFR inhibitor)^45, 46^ had little to no effect on NL09 replication but a more appreciable and sustained (5- to 100-fold) reduction, especially at the highest concentration used (0.5 μM) of NL11 replication. Defactinib (DF) (FAK/Pyk2 inhibitor)^47^ treatment resulted in robust reduction in viral titers (10- to 1,000-fold) in both NL09 and NL11 at the higher concentrations (2.5 and 5.0 μM). In NL09 infected cells, the reduction was more robust at earlier time-points (24 and 48 hpi), but in NL11 infected cells a larger reduction was observed at 24 and 72 hpi than at 48 hpi.

**Figure 2:**
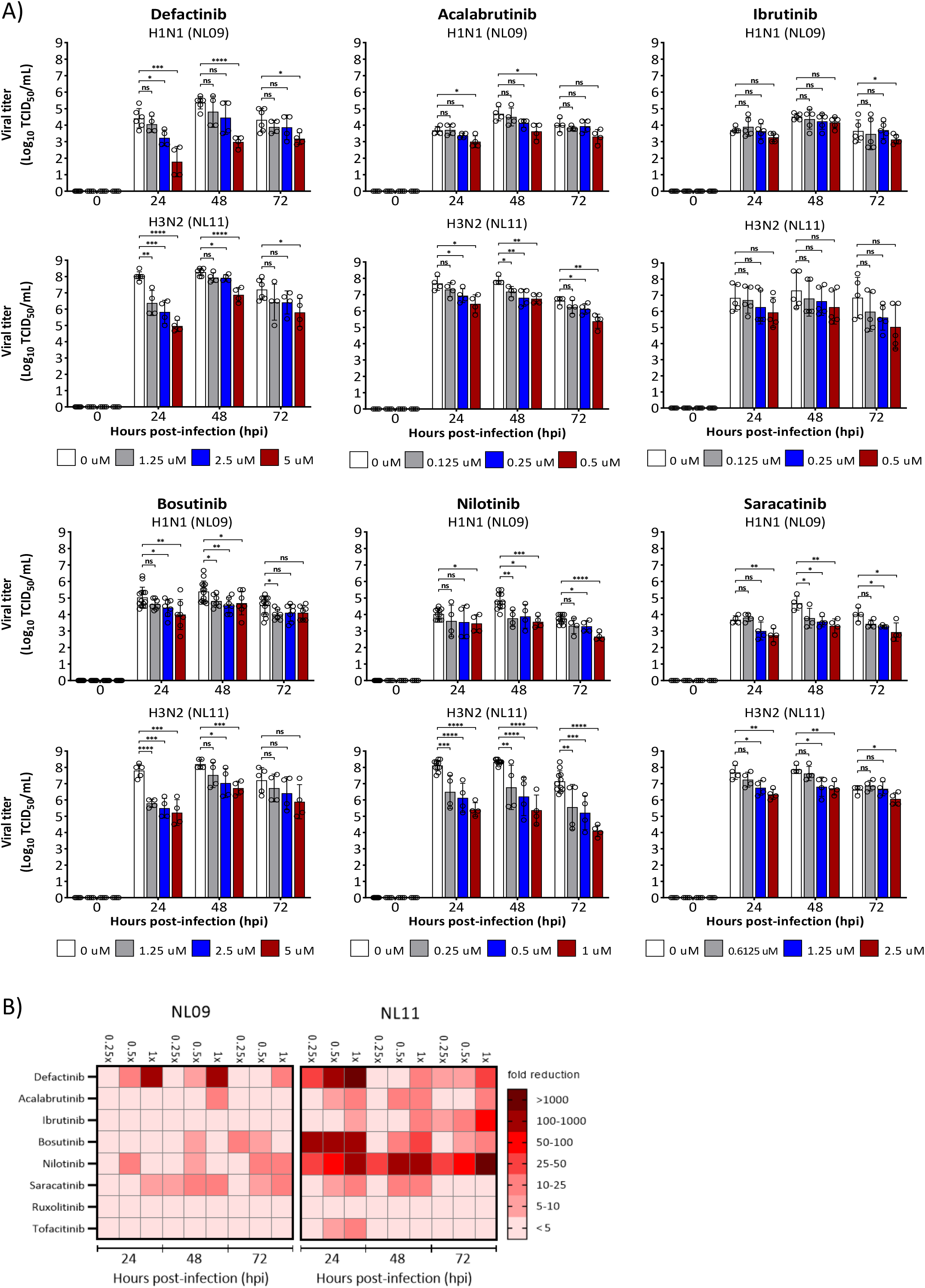
Effect of SMKI treatment on NL09 and NL11 replication at MOI 1. A549 cells were infected with pandemic H1N1 strain NL09 and seasonal H3N2 strain NL11 at MOI1 and incubated for 72h in the presence of SMKI concentrations of 0.25x, 0.5x and 1x with 1x being the highest non-toxic concentration under infection conditions (Tab. 1). At 24, 48, and 72 hpi, supernatants were collected and viral titers quantified by TCID_50_/ml assay (n = 4). Means ±SD are shown. *, P<0.05; **, P<0.01; ***, P<0.001; ****, P<0.0001; ns, not significant (P>0.05).

### NRTKIs affect IAV infectivity and cell viability during IAV infection

Host-targeted inhibitors can influence virus yield by affecting infectivity through effects on viral entry, RNA replication, increased host antiviral responses as well as impacting the viability of infected cells. To assess the effect of NRTKIs on viral infectivity and cellular viability, we used immunofluorescence microscopy to determine the number of infected cells as well as the total number of cells. A549 cells were infected with either NL09 or NL11 (MOI=1) in the presence of NRTKIs [0.5x]_max_, fixed at 48 hpi and stained to detect virus and nuclei (**Fig. 3A**). Surprisingly, treatment with most inhibitors resulted in a significant increase in infectivity despite the reduction in viral titers observed in **Fig. 2**. We observed robust increases in relative infectivity in cells treated with TF (NL09=217%, NL11=112%), RX (NL09=203%, NL11=116%) and DF (NL09=180%, NL11=139%). We only observed a marginal increase in relative infectivity (∼105%) in cells treated with either BO or NI in both NL09- and NL11-infected cells. We observed a marked decrease in relative infectivity in cells treated with AC (NL09=78%, NL11=74%) and SA (∼90% for both NL09 and NL11). Interestingly, we observed opposite effects on NL09 (increased)- and NL11 (decreased)-infected cells following treatment with IB (NL09=116% vs NL11=91%) (**Fig. 3B**).

**Figure 3.**
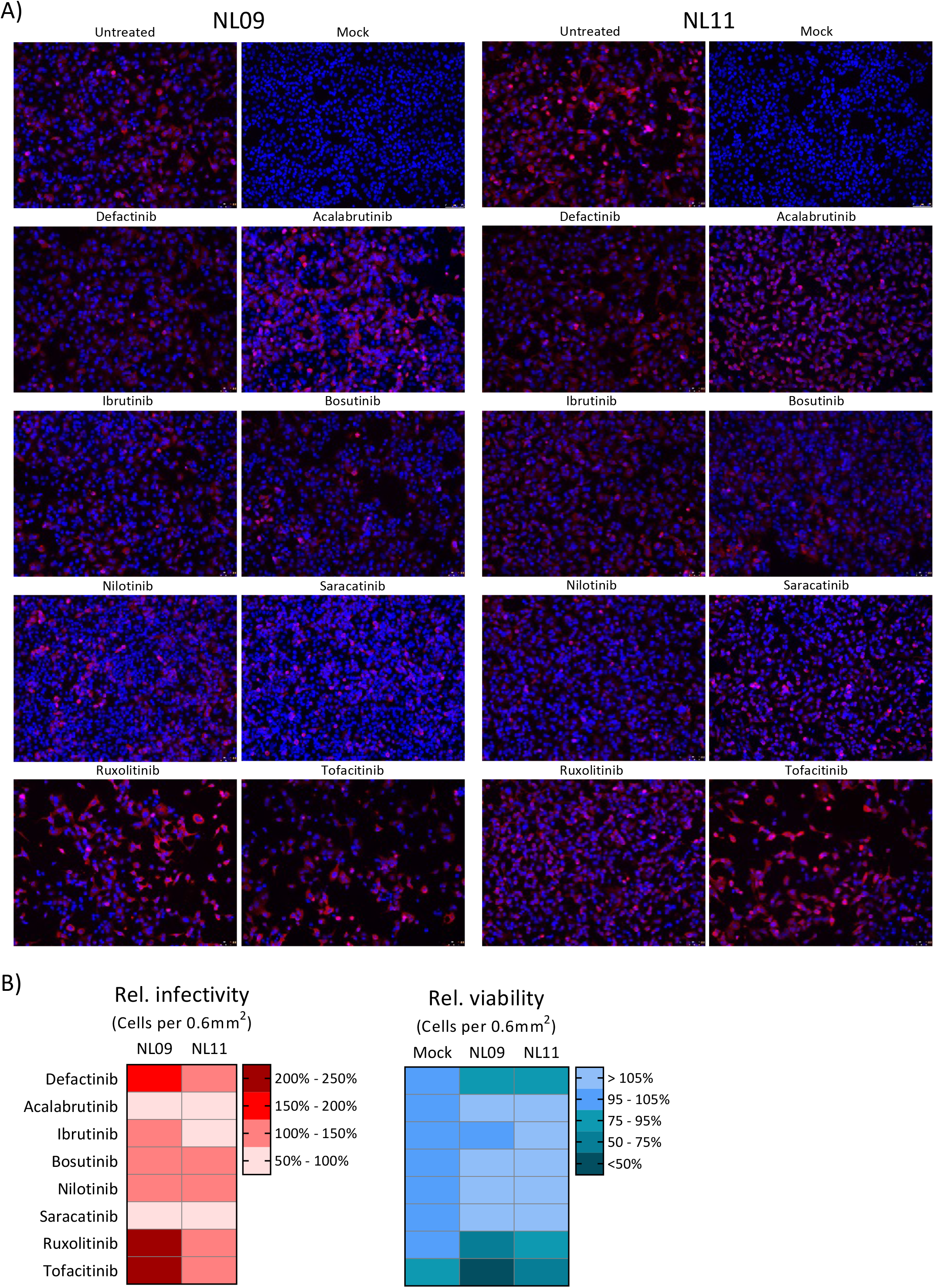
NRTKIs effects on cell viability and infectivity during infection. A549 cells were infected with pandemic H1N1 strain (NL09) and seasonal H3N2 strain (NL11) (MOI=1) and incubated in the presence of SMKIs (0.5x of the highest non-toxic concentration) for 48h. A) Fluorescence microscopy pictures were captured using a Leica DMi8 fluorescence microscope (representative field shown from an n=4/condition). Virus-infected cells were detected by anti-IAV NP antibody (red), and nuclei were detected using NucBlue Live ReadyProbes (blue). B) Heatmap visualization of NRTKIs affect cell viability (blue/green) and infectivity (red) relative to untreated infection in A549 cells. Images were quantified using ImageJ software suite. Data is based all fields represented in (A) (n=4/condition).

Next, we determined whether the reduction in titers was the result of reduced cell viability using the CellTiter Glo assay described above. Although at the NRTKI concentrations used [0.5x]_max_ we observed less than 5% cytotoxicity in mock-infected cells, IAV-infection led to synergistic cytotoxicity and decreased relative viability when treated with TF (NL09=40%, NL11=63%), RX (NL09=66%, NL11=89%) or DF (NL09=84%, NL11=92%). However, cell viability was increased by treatment of IAV-infected cells with AC (NL09=112%, NL11=109%), IB (NL09=102%, NL11=109%) BO (NL09=111%, NL11=114%), NI (NL09=110%, NL11=110%) or SA (NL09=150%, NL11=127%) (**Fig. 3B**). Taken together, our data suggest that decreased infectivity or cell viability alone do not account for the NRTKI induced reduction in viral titers we observed.

### The antiviral effect of NRTKIs is MOI-independent

Given the viability data and the limited reduction in viral titers above, we excluded RX and TF from further analyses and focused on the remaining six NRTKIs for further investigation. We assessed whether the inhibitory effects of NRTKIs we observed were dependent on the infectious dose used. We infected A549 cells with either a high MOI (MOI=3) or a low MOI (MOI=0.01) in the presence or absence of NRTKIs [0.5x]_max_. Culture supernatants were collected at 2, 24, 48, and 72 hpi and virus titers were determined by TCID_50_ assay. As we previously observed, the effect on the seasonal H3N2 (NL11) strain was more pronounced compared to that on the pandemic H1N1 (NL09) strain; presumably due to the faster growth kinetics of NL11 compared to NL09. However, similarly to cells infected at MOI=1 (**Fig. 2**), NRTKI treatment of cells infected at either high (3) or low (0.01) MOI resulted in viral titer reductions of at least 10-fold (1-log_10_) (**Fig. 4**). Treatment with either DF, BO or NI had a larger impact on early (24 hpi) viral replication, especially in NL11 infected cells. And although peak titers of untreated cells infected with NL11 were similar at either MOI, cells infected at MOI=0.1 exhibited the greatest reduction of ∼1,000-fold (3-log_10_). Although AC, IB and SA treatment had less of an impact on viral titers than DF, BO or NI, viral titer reductions were still up to 100-fold (2-log10) (**Fig. 4**).

**Figure 4.**
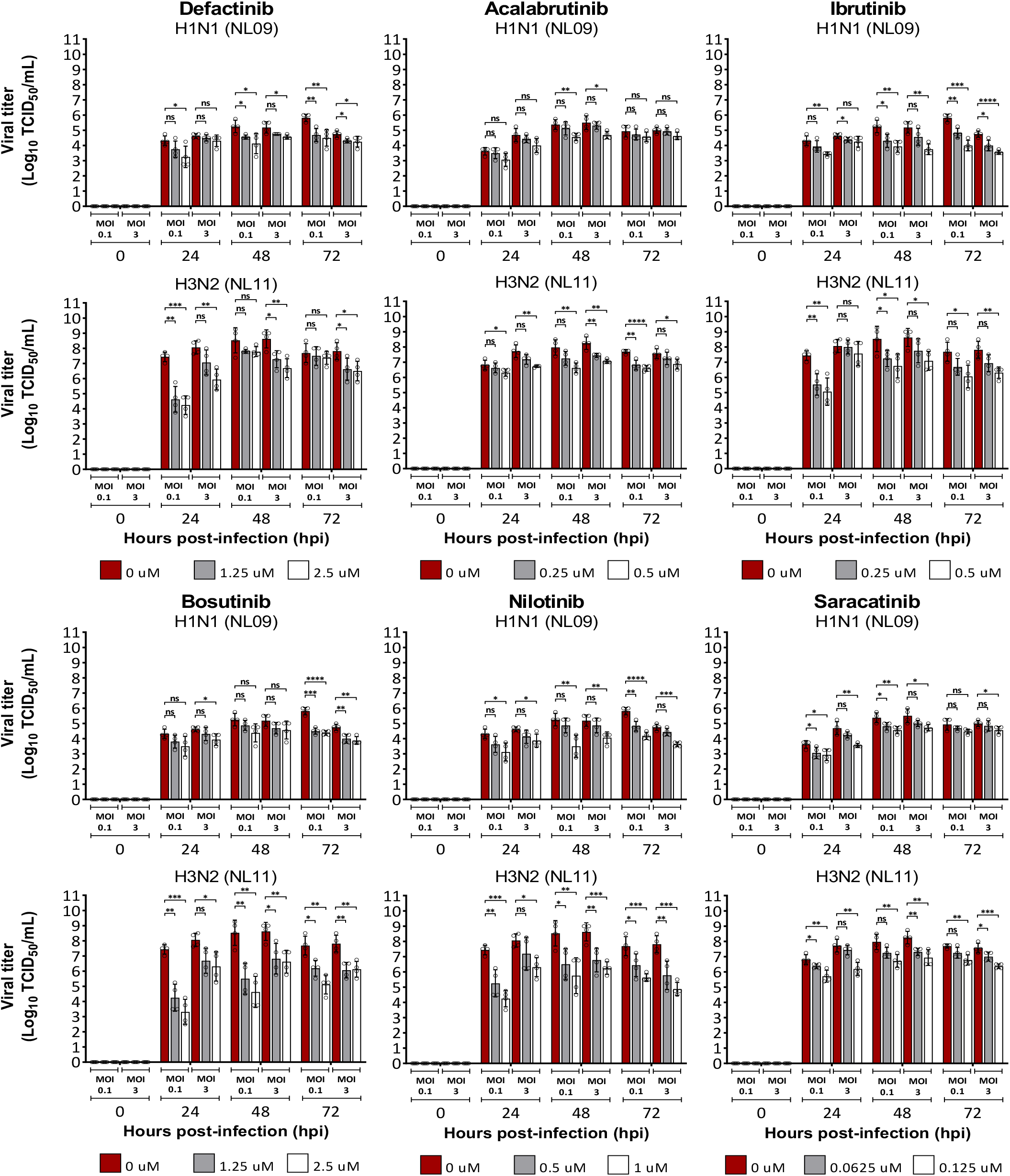
MOI-independent effect of NRTKIs on IAV infection. A549 cells were infected with pandemic H1N1 strain (NL09) and seasonal H3N2 strain (NL11) at either (MOI=0.1 or 3) and incubated for 72h in the presence of NRTKIs at 0.25x (gray) and 0.5x (white) of highest-toxic concentrations. At 24, 48, and 72 hpi, supernatants were collected and viral titers quantified by TCID_50_/ml assay (n=4). Means ±SD are shown. *, P<0.05; **, P<0.01; ***, P<0.001; ****, P<0.0001; ns, not significant (P>0.05).

### Human PCLS support robust IAV infection and confirm inhibitory effect of NRTKI treatment

We utilized human precision-cut lung slices (hPCLS) as a biologically relevant *ex vivo* model that more faithfully represents lung tissues than either 2D monolayer cultures or 3D air-liquid-interface (ALI) cultures ^33, 34^. Following hPCLS preparation, they were in culture for up to 4 weeks; no gross alterations in cell type or morphology were observed and cilial beating was observed in all used hPCLS (8 donors; n=24). We first sought to identify an infectious dose to synchronize peak titers in hPCLS infected with either NL09 or NL11 as these two strains have different replication kinetics *in vitro*. hPCLS were infected with 10^4^, 10^5^, or 10^6^ TCID_50_ / 200 µL with either NL09 or NL11. To limit donor-heterogeneity effects, we used hPCLS from 8 donors (n=24/virus). Culture supernatants were collected and replenished at 2, 16, 24, 48, 72, 96, and 144 hpi and virus titers were determined by TCID_50_ assay using the collected supernatants. Interestingly, the 10^6^ dose yielded maximal titers that were lower than the other doses and were achieved by 24 hpi with either strain. However, the highest peak titers were achieved at 48 hpi following infection with either 10^4^ or 10^5^ doses of NL11, but only in the 10^5^ dose of NL09 (**Fig. 5A**). Based on these results the 10^5^ dose was used in all our subsequent hPCLS infections.

**Figure 5.**
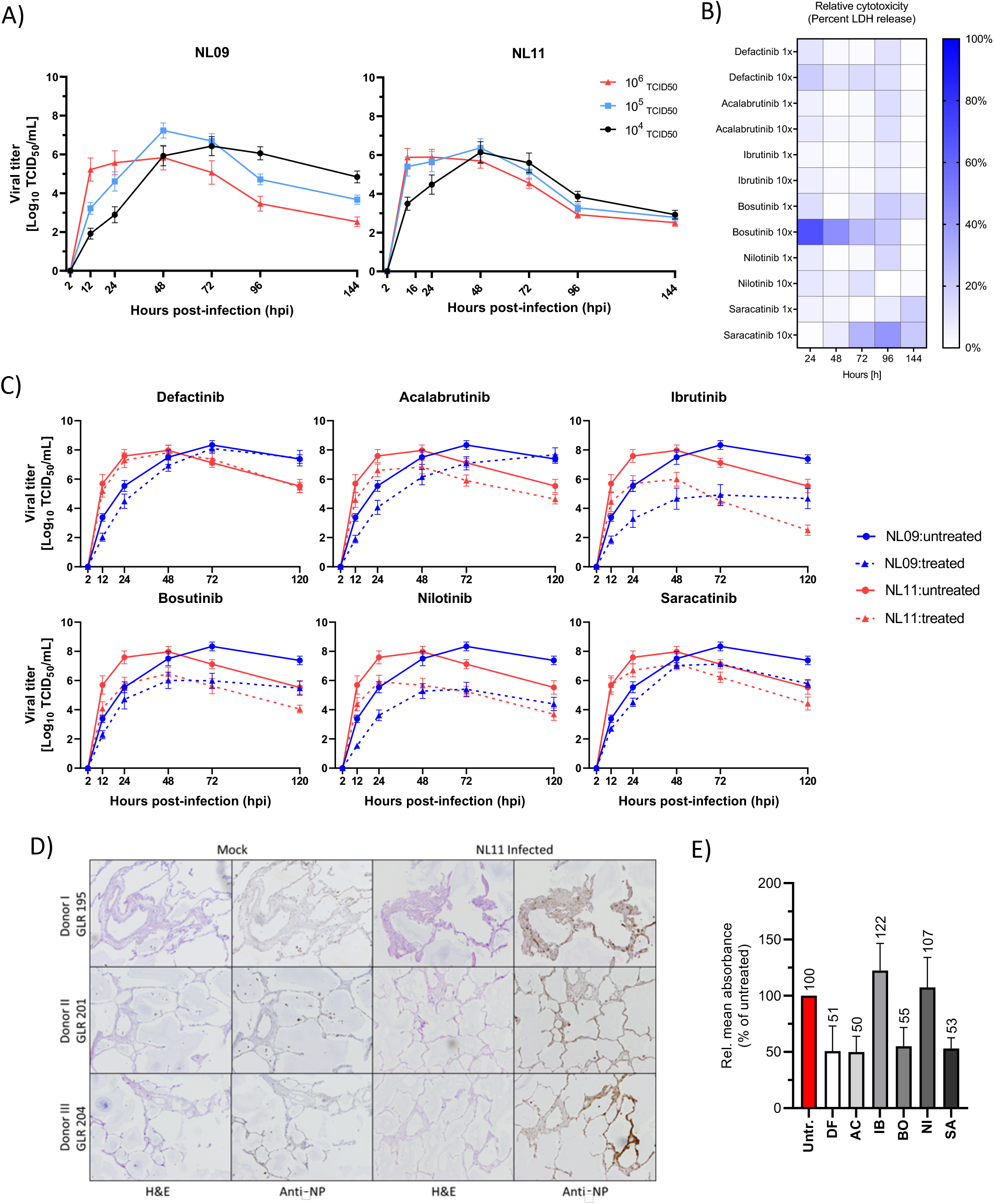
Effect of NRTKI treatment on *ex vivo* IAV infection. A) hPCLS were infected with pandemic H1N1 strain (NL09) and seasonal H3N2 strain (NL11) at increasing doses (10^4^, 10^5^ and 10^6^ TCID_50_/200 ul). At 2, 16, 24, 48, 72, 96, and 144 hpi culture supernatants were sampled and viral titers quantified by TCID_50_/ml assay. The supernatants were replenished after collection at every time point (8 donors; n=24/virus); means ±SEM are shown. B) Heatmap visualization of NRTKI cytotoxicity based on LDH release of hPCLS normalized to DMSO control, and relative to 1% Triton-X 100 treated cells; NRTKIs were treated with 1x and 10x of highest cytotoxic concentration on A549 cells (Fig. 1) for up to 144h. At every time point, LDH release was determined using LDH-Glo Cell Viability Assay (8 donors / n=24). C) PCLS were infected with NL09 or NL11 (10^5^ TCID_50_ / 200 ul) and incubated for 120h in the presence of NRTKIs (Defactinib 50uM; Acalabrutinib 5uM; Ibrutinib 5uM; Bosutinib 5uM; Nilotinib 10uM; Saracatinib 0.125uM). Growth curves from supernatants collected at 2, 12, 24, 48, 72, and 120 hpi were quantified by TCID_50_/ml assay (3 donors; n=6/condition); means ±SEM are shown. D) NL11 infected hPCLS were fixed 120 hpi and the PFA-fixed paraffin-embedded (PFPE) PCLS were were cut into 2 µm thick section. Shown is H&E staining and viral anti-NP immunohistochemical staining (brown). Original magnification 10x. E) Semi-quantitative analysis of virus infected (anti-NP staining) was performed for all tested NRTKIs using FIJI image-analysis software platform and normalized to whole section areas.

Next, we determined the tolerability of hPCLS to our NRTKIs candidates. hPCLS were treated with either the [1x or 10x]_max_ (1x is highest non-toxic NRTKI concentration defined using A549 cells). Culture supernatants were collected and replenished at 24, 48, 72, 96, and 144 h. Lactate dehydrogenase (LDH) release into the culture supernatant due to loss of plasma membrane integrity is an indicator for cytoxicity. Using a bioluminescence based LDH detection assay (LDH-Glo Cytotoxicity Assay), we quantified LDH released into the collected supernatants following NRTKI treatment. As a positive control for cytotoxicity, hPCLS were treated with 0.1% Triton-X 100; DMSO-treated hPCLS were used as a vehicle control (**Fig. 5B**). Our cytotoxicity cut-off was 20% of the positive control treatment; none of the NRTKIs surpassed this cut-off at [1x]max. However, [10x]_max_ concentrations of DF (50 μM), BO (50 μM) and SA (1.25 μM) showed significantly higher cytotoxicity that was above the relative 20%-cutoff; therefore, only the 1x concentrations of these NRTKIs were used in subsequent hPCLS experiments.

### NRTKIs inhibit *ex vivo* IAV infections

Based on our *in vitro* data, we tested six NRTKIs (AC, BO, DF, IB, NI and SA) for their antiviral potential. hPCLS from 3 donors (n=6/virus/condition) were infected with 10^5^ TCID_50_ of either NL09 or NL11 and then treated with NRTKIs (DF 5μM; AC 5μM; IB 5μM; BO 5μM; NI 10μM; SA 0.125μM). Culture supernatants were collected and replenished from the same wells at 2, 12, 24, 48, 72, and 120 hpi and virus titers were determined by TCID_50_ assay. We observed a significant and robust reduction in viral titers of at least 10-fold or 1-log_10_ (DF treatment) to more than 1,000-fold or 3-log10 (IB and NI treatments) using all the NRTKIs (**Fig. 5C**). Moreover, unlike what we observed in A549 cells, NRTKI-mediated IAV inhibition was observed as early as 12 hpi and maintained at 120 hpi; beyond the times when peak titers were achieved (48-72 hpi) (**Fig. 5C**).

Next, we assessed the effect of NRTKI treatment on viral spread and associated damage to the epithelium. Considering we did not observe significant differences in titer reductions between NL09 and NL11, only data for NL11 is shown (**Fig. 5D**). At 120 hpi, mock- and IAV-infected hPCLS (n=3/virus/condition) were fixed and paraffin-embedded. Tissue sections (2 μm thick) were cut, stained and immunoprobed. H&E staining indicated that no gross alterations in cell composition or the epithelium were observed in mock-infected cells indicating viability of hPCLS. Typical morphological changes associated with IAV infections were observed. Using consecutive sections from those H&E stained, we detected the viral spread using antibodies to IAV-NP (**Fig. 5D**). We observed specific staining of viral antigen in all observed cell-types including type I/II pneumocytes and endothelial cells. Additionally, staining intensity and quantity was reduced in NRTKI treated hPCLS compared to untreated hPCLS.

We carried out semiquantitative analysis of the acquired images to compare the effect of the inhibitors on viral spread in hPCLS with variable tissue density. IAV-NP signal was normalized to H&E staining of the respective regions in consecutive tissue section. Treatment with DF, AC, BO and SA significantly reduced the infectivity by > 50%, whereas treatment with IB and NI had limited effects on infectivity (**Fig. 5E**).

### Stability of NRTKI inhibition

Host-directed antivirals/therapeutics are thought to have a higher barrier of resistance than their virus-targeted counterparts. To determine the stability of the antiviral effect of our NRTKIs, we phenotypically assessed the possibility of the emergence of resistant escape variants ^48, 49^. We passaged both NL09 and NL11 viruses (MOI=0.001) in the presence of NRTKIs [1x]_max_ in MDCK cells for 5 passages. Untreated virus stocks were also passaged as a control. At each passage, virus supernatant was quantified and used to inoculate the next passage at MOI=0.001 again. In untreated passages, the virus titers for both NL09 and NL11 was similar from passages 1 to 5 but were significantly lower in all treated passages (**Fig. 6A**); the reduction was comparable to what we originally observed in A549 cells. The viral titers were stable at all passages indicating that no resistance mutations were acquired. To rule out the possibility that the NRTKIs directly interact with the virion and inhibit its attachment or entry, we pre-incubated virus stocks with NRTKIs [1x]_max_ for 2 h then diluted the virus stocks 1:1000 to minimize the effects on host cells, and infected A549 cells. At 72 hpi, culture supernatants were collected and virus titrated by TCID_50_ assay. As expected, pre-treatment of virus with NRTKIs had no effect on viral titers, indicating that the observed effects are due to host-cell effects (**Fig. 6B**).

**Figure 6.**
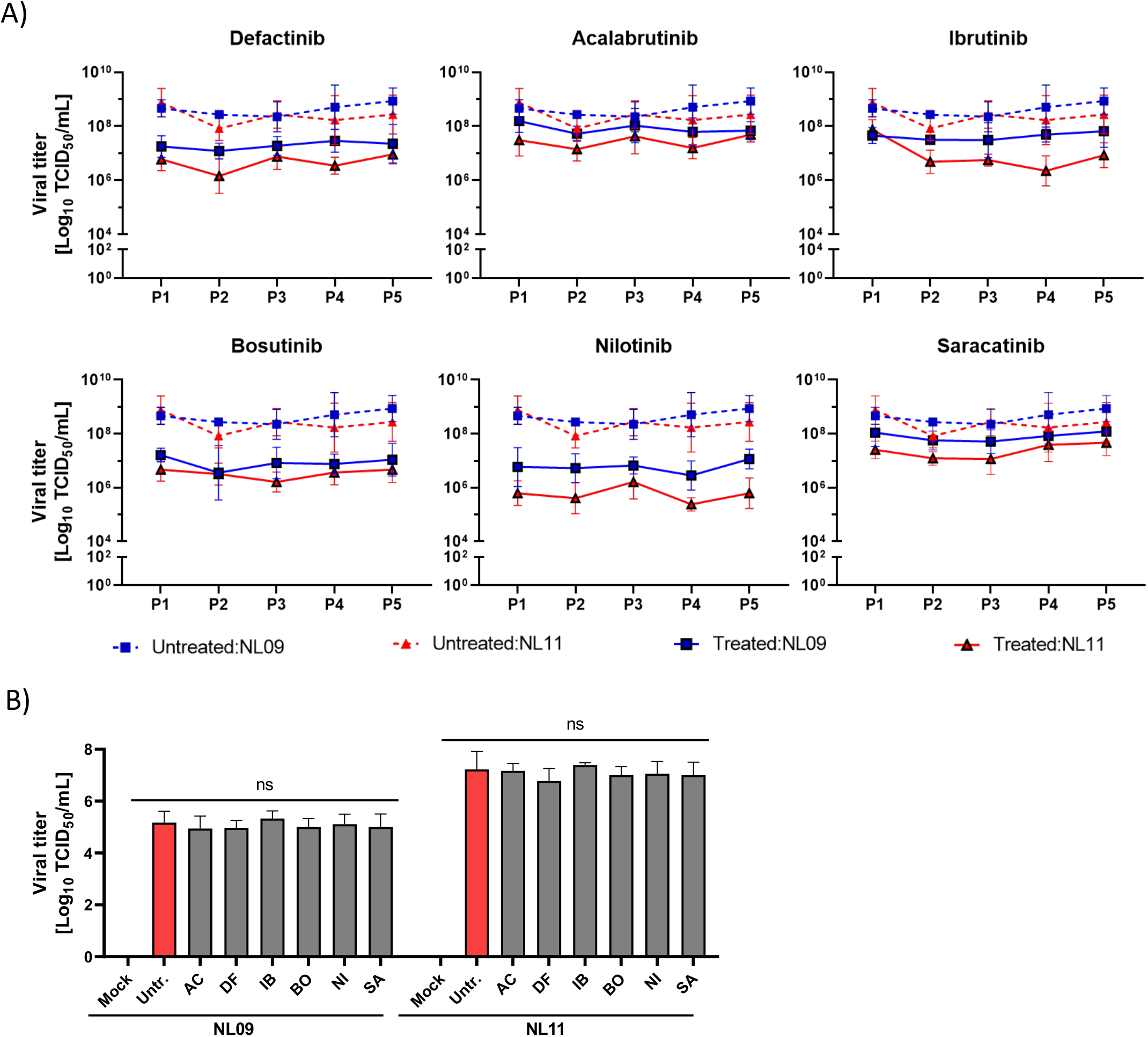
Stability of NRTKI treatment effect. (A) Effect of SMKI treatment on NL09 and NL11 replication at MOI=0.001 over serial passages was determined by growth curves using MDCK cells infected with pandemic H1N1 strain (NL09) or seasonal H3N2 strain (NL11) at MOI=0.001 and incubated for 72h in the presence of the highest non-toxic NRTKI concentrations (n=2). At 72 hpi, samples were collected, and viral titers were assessed by TCID_50_/ml assay on MDCK cells (n = 4). MDCK cells were infected as before at MOI=0.001 according to the assessed viral titers. Means ±SD are shown. (B) NL09 and NL11 virus stocks were pre-incubated with control (DMSO) or the 1x concentration of the respective NRTKI for 4h at 37°C. A549 cells were then infected using a 1:1000 dilution of the NRTKI pre-incubated virus stocks and incubated for 72h. At 72 hpi, samples were collected, and viral titers were assessed by TCID_50_/ml assay (n=3). Means ±SD are shown. *, P<0.05; **, P<0.01; ***, P<0.001; ****, P<0.0001; ns, not significant (P>0.05).

### Selected NRTKIs inhibit viral entry

Kinases regulate every step of the infection cycle and a single kinase can affect multiple steps ^28, 29^. We first assessed whether our NRTKIs impaired viral entry. A549 cells were pretreated for 2 h, then chilled on ice for 15 min and infected at a high MOI (MOI=10) on ice for 30 min to synchronize the infection and allow binding of the virus to cell surface receptors but not trafficking of virions. Unbound and non-internalized virus was washed away with room temperature PBS. Cells were then incubated with prewarmed infection media in the presence or absence of NRTKIs. At 0.5 hpi, cells were fixed, stained to detect viral NP, F-actin and nuclei and analyzed by confocal microscopy. We observed significant retention at the membrane and periphery of the cell following DF and IB treatment (**Fig. 7**). Surprisingly, no virus was detected in response to BO treatment and the F-actin network was not detectable. Given the sustained viability of BO-treated cells, it is likely that the altered actin dynamics are well tolerated. No significant changes were detectable in AC, NI or SA treated cells suggesting these inhibitors did not affect viral entry under our tested conditions (**Fig. 7**).

**Figure 7.**
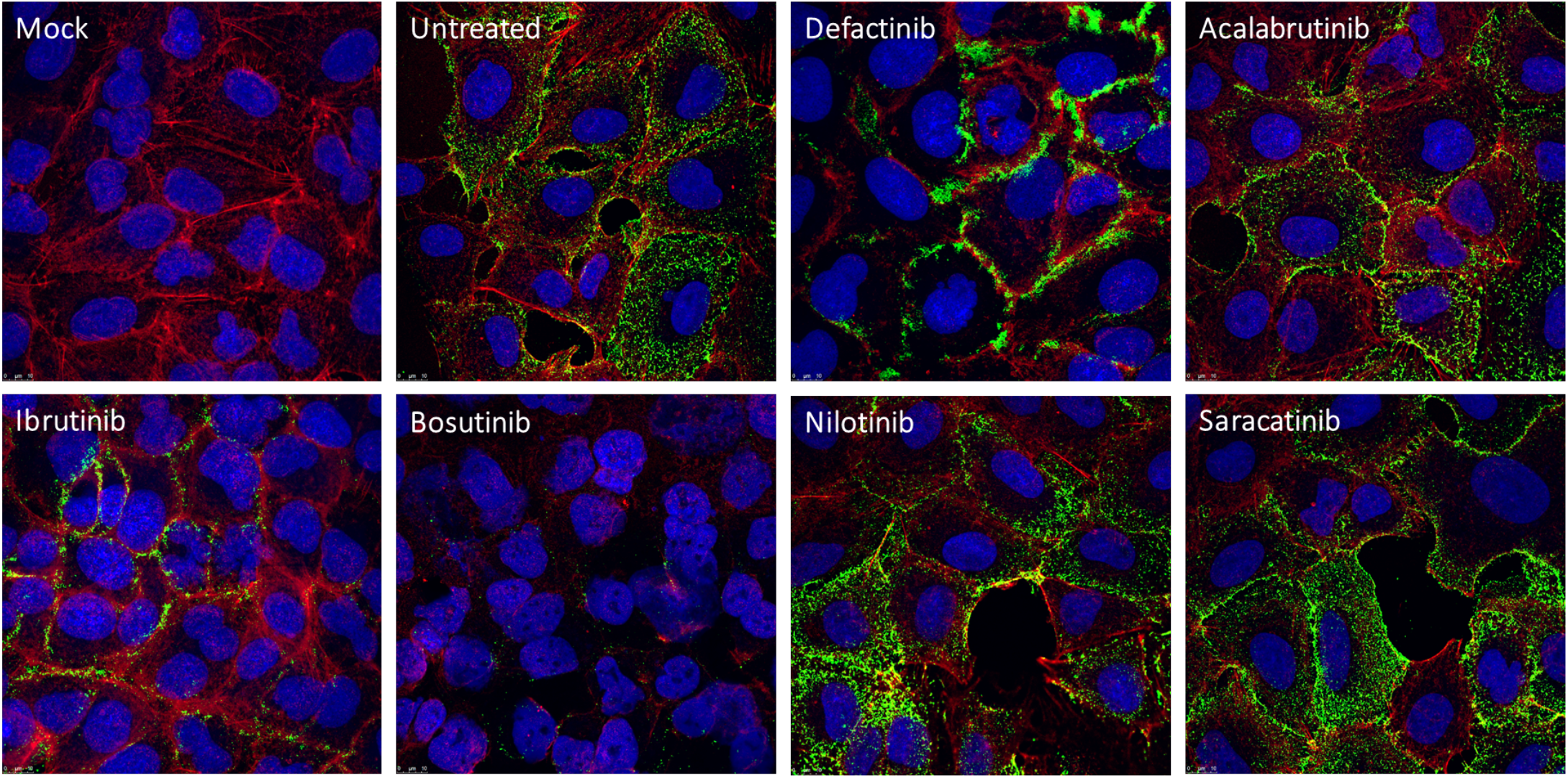
The effect of NRTKI on viral entry. A549 cells were pretreated with NRTKIs for 2h and then infected with the NL09 and NL11 strains (MOI=10) for 0.5h in the presence or absence of the inhibitors. The cells were fixed and permeabilized. Virions were detected by anti-NP (green) antibody, F-Actin was detected by ActinRed-555 (red), and nuclei were detected using NucBlue Live ReadyProbes (blue). and virion localization was assessed by confocal microscopy using a 63x oil immersion objective (n = 2).

### NRTKIs exert differential effects on IAV polymerase activity

Next, we assessed the effect of our NRTKIs on viral RNA replication using the pPOLI-358-FFLuc reporter plasmid, which encodes a firefly luciferase gene under control of the viral nucleoprotein (NP) promoter. In this system, luciferase activity is a surrogate for viral polymerase activity ^50–52^. A549 cells were transfected with pPOLI-358-FFluc and pmaxGFP plasmids (transfection control). At 24 hpt, cells were either infected with NL09 or NL11 at MOI=1 in the presence or absence of indicated NRTKIs at either [0.5x or 1x]_max_. At 48 h post-transfection (hpt) (∼24 hpi), luciferase activity was measured, normalized to GFP expression (MFI) and represented as relative of untreated infected cells. We observed a significant reduction in polymerase reporter activity in response to AC (NL11 only), IB (NL09 only), BO, NI and SA (NL11 only) (**Fig. 8A, left panel**). Although the magnitude of reduction was higher in NL09-infected than NL11-infected cells, a significant reduction was more readily observed in NL11 infected cells at a lower NRTKI concentration compared to NL09 infected cells. DF also reduced reporter activity (NL09=20%, NL11=13%), however this reduction was not statistically significant. At 24 hpi, a 3-fold increase in reporter activity could be observed in untreated NL11-infected cells over untreated NL09-infected cells; this is in-line with faster replication kinetics of NL11 compared to NL09 (**Fig. 8A, right panel**).

**Figure 8.**
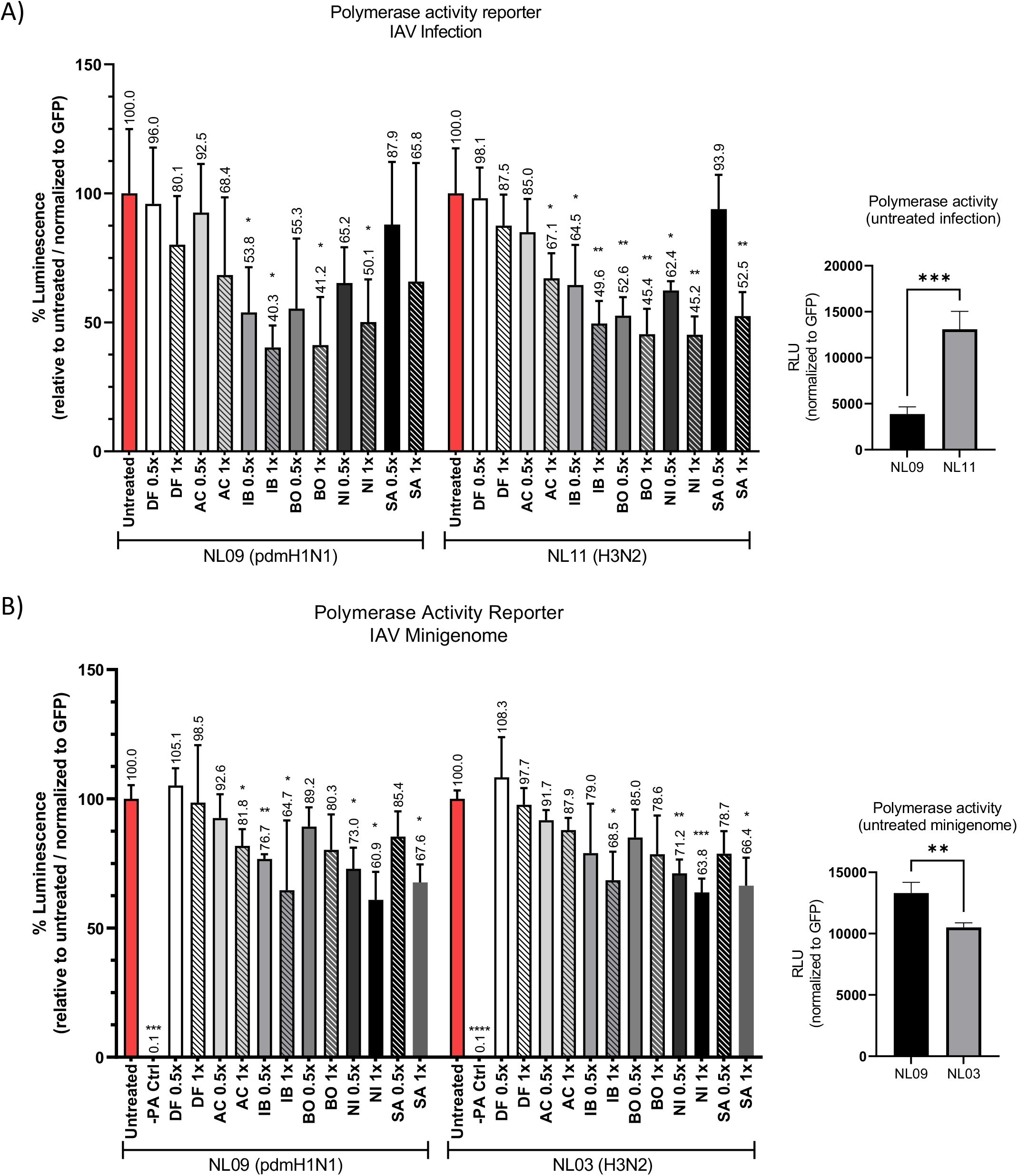
NRTKIs effects on IAV RNA replication. (A) A549 cells were transfected with pPOLI-358-FFluc and pmaxGFP plasmids. At 24 hpt, cells were either infected with NL09 or NL11 at MOI=1 in the presence or absence of indicated NRTKIs. At 48 hpt (24 hpi), luciferase activity was measured and normalized to GFP expression (MFI). (B) A549 cells were transfected with pPOLI-358-FFluc and pmaxGFP plasmids and co-transfected with either NL09 or NL03-minigenome plasmids. At 6 hpt, the indicated NRTKIs were added to the medium. At 30 hpt (24 h of treatment), luciferase activity was measured and normalized to GFP MFI. Bars indicate values relative to infected untreated cells normalized to GFP. For each system luminescence of untreated infected A549 cells relative to GFP displays the replication kinetics of the respective polymerase complex. All measurements were taken in triplicates from triplicate samples (n=3). Error bars indicate ± standard deviation (SD). *, P<0.05; **, P<0.01; ***, P<0.001; ****, P<0.0001; ns, not significant (P>0.05). P-values determined by Brown-Forsythe and Welsh ANOVA compared to untreated.

To better dissect the direct effect on viral RNA replication in the absence of confounding factors due to NRTKI influences on viral entry and host responses, we utilized an established minigenome system. A549 cells were transfected with pPOLI-358-Ffluc and pmaxGFP plasmids and co-transfected with either NL09 or NL03-minigenome plasmids that encode the viral NP, PA, PB1 and PB2 replication complex proteins. At 6 hpt, the indicated NRTKIs were added to the medium at either [0.5x or 1x]_max_. At 30 hpt (24 h of treatment), luciferase activity was measured, normalized and represented as above. In this context, AC (NL09 only), IB, NI and SA treatments significantly reduced polymerase activity (**Fig. 8B**). In contrast to what we observed in infected cells, the magnitude of reduction in polymerase activity was comparable in NL09- and NL11-minigenome transfected cells. Interestingly, polymerase activity was significantly higher in untreated H1N1 (NL09) than H3N2 (NL03) minigenome-transfected cells.

### NRTKIs do not affect innate immune responses during IAV infections

Given that our NRTKIs reduced viral titers by affecting either viral entry or replication, we speculate whether these effects were coupled to altered innate immune signaling. Activation of STAT3 by IFN type-I, -II, and -III as well as by interleukins results in phosphorylation of STAT3 at Y705. We assessed the level of pSTAT3 (pY705) in A549 cells infected with either NL09 or NL11 (MOI=1) in the presence or absence of NRTKIs at [1x]_max_. At 18 and 48 hpi, total proteins were isolated from whole cell lysates, separated by SDS-PAGE and analyzed by immunoblotting. We observed a decrease in the relative pSTAT3/total STAT3 ratio in untreated NL09-infected cells at 18 and 48 hpi (**Fig. 9A**) compared to mock-infected cells (18 h = 159%, 48 h= 225%). This reduction was less striking in NL11-infected cells (**Fig. 9B**). Only DF treatment resulted in a significant reduction of pSTAT3 relative to untreated infected cells (NL09 = 5% to 8%, NL11 = 5% to 14% of untreated). None of the other NRTKIs showed a significant effect.

**Figure 9.**
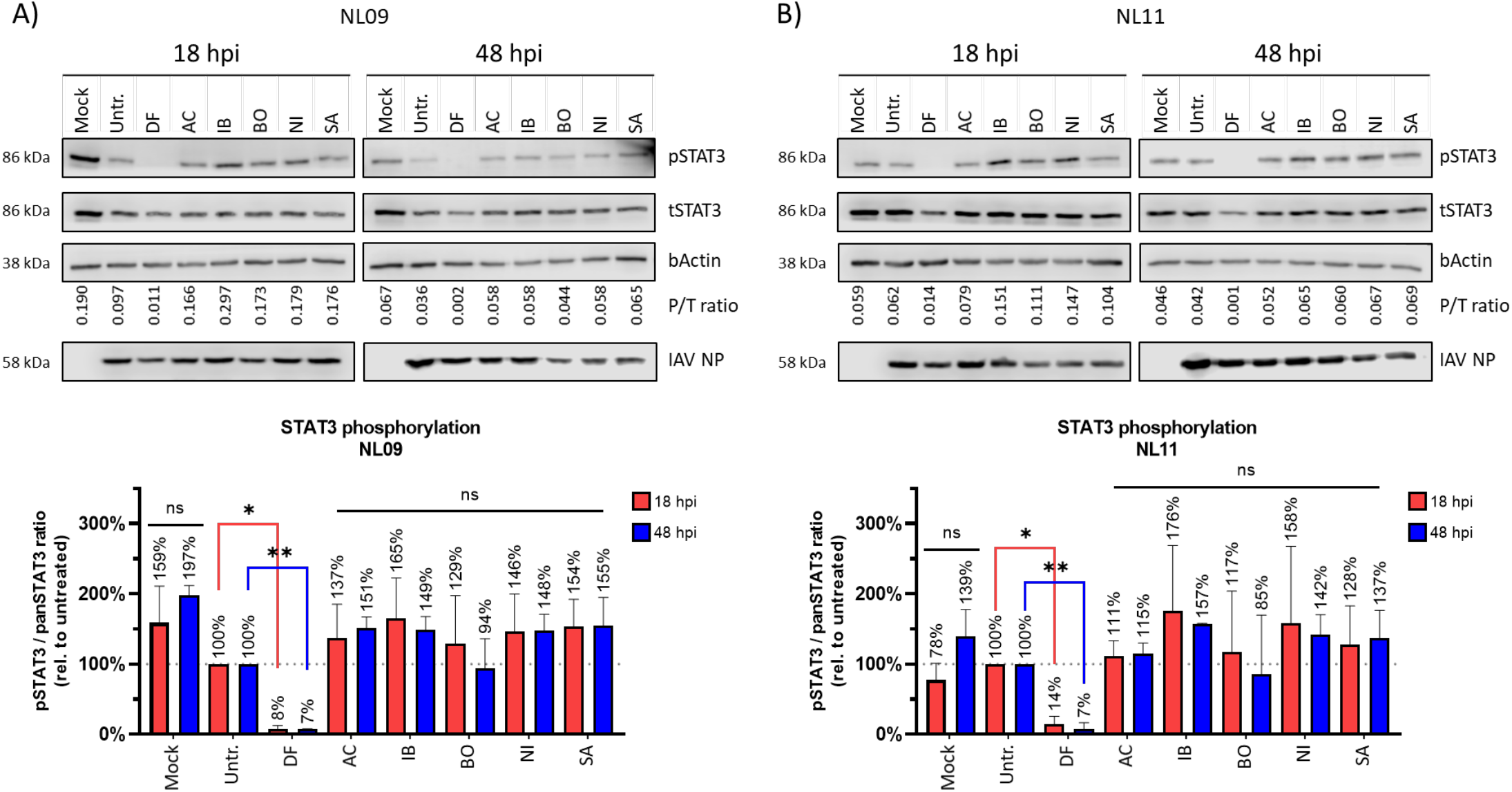
Effect of NRTKIs treatment on STAT3 activation. A549 cells were infected with NL09 (A) or NL11 (B) at MOI=1. Total proteins were isolated from whole cell lysate at 18 and 48 hpi and immunoblot assay was performed for phospho- and total STAT3, IAV-NP and bActin. Chemiluminescence was detected and quantified using the Li-Cor C-DiGit and Image Studio 5.1 CLX software. All measurements were taken from two western blots from two independent experiments (n=2). All values are relative to untreated virus-infected cells. P=phosphor, T=total. Error bars indicate ± standard deviation (SD). *, P<0.05; **, P<0.01; ***, P<0.001; ****, P<0.0001; ns, not significant (P>0.05). P-values determined by students t-test compared to untreated virus-infected cells.

Phosphorylation of NFkB p65 (pNFkB) at S536 results in activation of the NFkB pathway. We did not detect an increase in the relative pNFkB/total NFkB ration in NL09-infected cells compared to mock-infected cells at either 18 or 48 hpi following treatment with any of the NRTKIs (**Fig. 10A**). In contrast, we observed a slight increase in the pNFkB/total NFkB ratio in untreated NL11-infected cells at 18 and 48 hpi (**Fig. 10B**) compared to mock-infected cells (18 h = 75%, 48 h= 89%). Although we did observe a reduction in NFkB activation following DF treatment of NL11-infected cells (18 h = 64%, 48 h = 68% of untreated), this reduction was not statistically significant. We next confirmed that NFkB signaling is not impaired in our system. Mock-infected cells were treated with high-molecular weight poly(IC), a synthetic dsRNA, at either 50 or 200 ng/ml. Following infection of cells with NL09 or NL11 (MOI=1), cells were treated with 200 ng/ml poly(IC). At 18 and 48 hpi, total proteins were isolated from whole cell lysate, separated by SDS-PAGE and analyzed by immunoblotting. At 18 hpi, the relative pNFkB/total-NFkB ratio significantly increased in response to poly(IC) stimulation with both low and high concentrations (∼7-fold relative to untreated 18 h mock) (**Fig. 10C**). Similarly, poly(IC) treatment of NL09 or NL11 infected cells had a robust increase in NFkB activation (NL09 = ∼10-fold, NL11 = ∼17-fold relative to untreated 18 h mock) (**Fig. 10C**). By 48 hpi, the observed NFkB activation was back to untreated-levels in mock, NL09 and NL11 infected cells. These data further support our results that IAV-induced NFkB activation is limited later during infection; likely suppressed by the viral NS1 (**Fig. 10C**).

**Figure 10.**
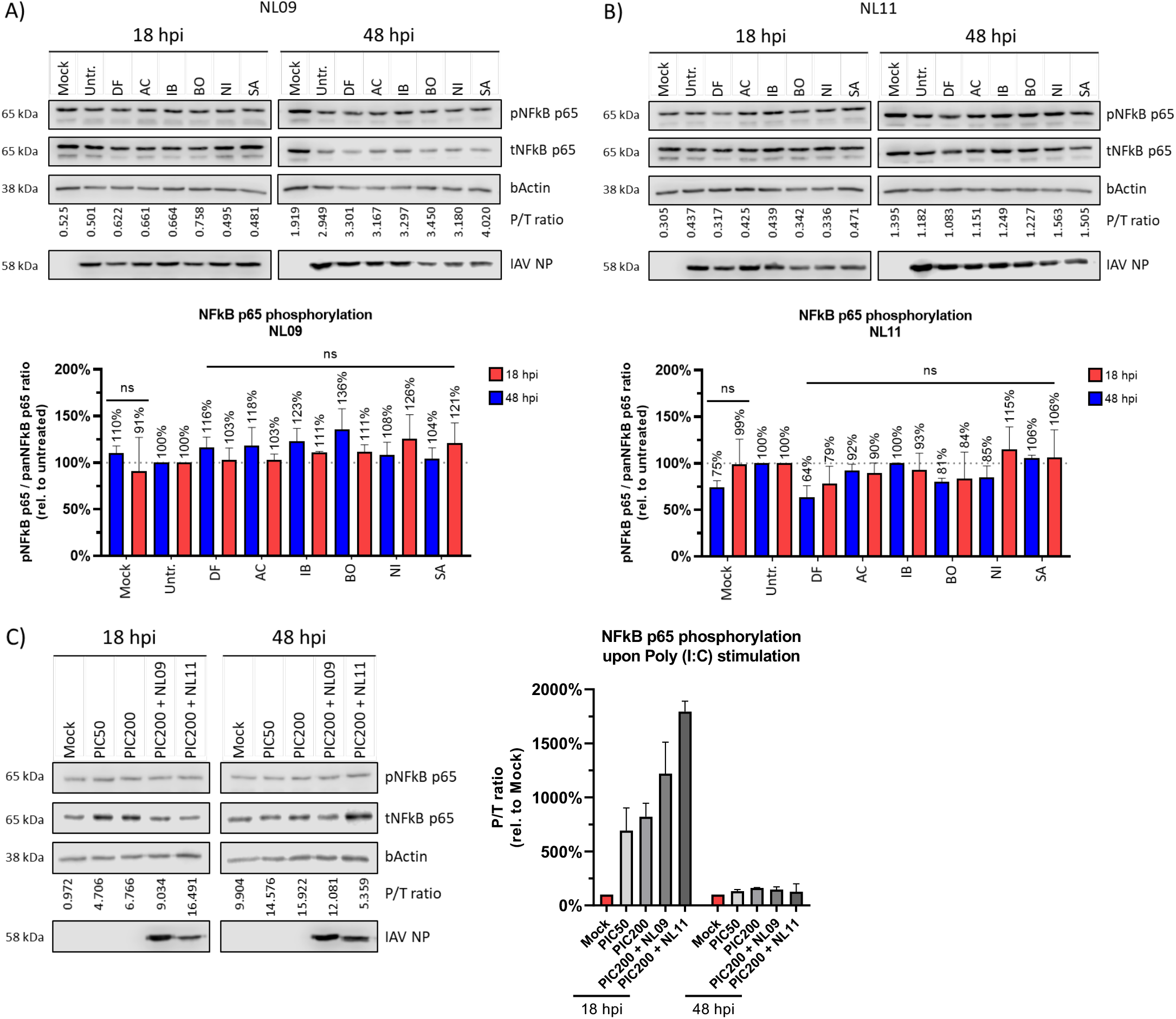
Effect of NRTKIs on NFkB activation. A549 cells were infected with NL09 (A) or NL11 (B) at MOI=1. Protein was isolated from whole cell lysate at 18 and 48 hpi and immunoblot assay was performed for pNFkB p65, panNFkB p65, IAV NP and bActin. Detected signal was quantified using the Li-Cor C-DiGit and Image Studio 5.1 CLX software. (C) NL09, NL11 or mock infected cells were treated with poly(IC) at concentrations of 50 ng/ml and 200 ng/ml. Total proteins were isolated from whole cell lysate at 18 and 48 hpi and immunoblot assay was performed for phospho- and total NFkBp65, IAV NP and bActin. All measurements were taken from two western blots from two independent experiments (n=2). All values are normalized to bActin and relative to untreated of the respective time-point. Error bars indicate ± standard deviation (SD). P=phosphor<u>y</u>lated and T=total. *, P<0.05; **, P<0.01; ***, P<0.001; ****, P<0.0001; ns, not significant (P>0.05). P-values determined by students t-test compared to untreated virus infected cells.

## DISCUSSION

Despite their clear susceptibility to rapidly arising resistance mutations, virus-targeted antivirals are still the only available class of antivirals against respiratory viruses such as influenza. In this study, we screened FDA approved SMKIs currently in clinical use against cancers and autoimmune diseases for their antiviral potential against IAV infections. Six of the eight NRTKIs we tested showed a potent inhibition of pandemic A(H1N1)pdm09 and seasonal A(H3N2) IAV strains with little to no impact on cell viability *in vitro*. We further validated these NRTKI candidates using a faithful *ex vivo* model of human PCLS. We identified the step(s) of the viral replication cycle affected by each compound. In doing so, we provide valuable information on the interplay of signaling pathways regulating these steps and the likely kinases involved.

Kinase dysfunction often leads to malignancies and tumor immune evasion. Therefore, kinases have been heavily targeted for cancer treatments using selective inhibitors ^26^. At the molecular level, most kinases regulate signaling pathways via catalytic and/or protein-scaffolding activities. Phosphorylation often triggers changes in protein conformation, enzymatic activity or subcellular localization; all of which allow fine-tuning of protein-protein interactions to mediate stimuli-specific responses ^53^. Indeed, host kinases play a critical role in IAV entry, replication, and release as well as viral evasion/suppression of hosts immune responses. These processes often require phosphorylation of viral proteins by mostly unidentified kinases ^54–65^.

We carried out our initial screening using A549 cells (ATII lung adenocarcinoma); however, due to potential biases associated with aberrant expression and kinase activity of cancerous cell-lines, we validated candidate NRTKIs using human PCLS (hPCLS). Unlike 2D monolayers or 3D well-differentiated air-liquid interface (ALI) cultures, PCLS preserve the native lung tissue architecture, cellular composition including endothelial, ATI and ATII epithelial cells, fibroblasts and maintain the native extracellular matrix ^33–35^. Moreover, the tissue tropism and infectivity of certain viruses may not be accurately represented *in vitro* due to the absence of relevant cell-cell interactions that can influence infectibility and host responses ^66^. Accordingly, we observed a similar discrepancy in which the strain-dependent differences observed in NRTKI-treated A549 cells were not observed in hPCLS, suggesting that the variations between IAV strains in A549 might be an *in vitro* artifact. Moreover, we observed a wide tissue tropism in hPCLS which suggests that while ATII cells may support more efficient infection, other cell-types of the lung are readily infectible as well. Nevertheless, we observed robust viral titer reductions in both systems following NRTKI treatment that were not readily explained by a reduction in infectivity or cell viability. Considering that the smallest reductions in viral titers were observed following Defactinib (DF) < Acalabrutinib (AC) < Saracatinib (SA) < Bosutinib (BO) treatment, hPCLS infectivity was reduced by ∼50% by each of those NRTKIs. Similarly, Ibrutinib (IB) and Nilotinib (NI) had the largest effect on viral titers but increased infectivity (NI:∼22%, IB=∼7%). Together our data suggest that reduction in infectivity of either A549 cells or hPCLS does not fully account for the potent reduction in viral titers and supports a *bona fide* effect of NRTKIs on either viral entry or RNA replication.

Although there is a growing body of *in vitro* and *in vivo* evidence to support the use of kinase inhibitors, not a single SMKI has been approved or licensed for the treatment of influenza virus infection so far ^4, 13, 14, 17, 60^. SMKI selectivity remains a contentious topic and has been a hurdle to the pursuit of kinase inhibitors as antivirals. Assumptions made regarding “off-target effects” are attributed to changes in phosphorylation or activation of proteins/pathways besides the intended target. While compounds still in the pre-clinical development phase require target validation, the selectivity of compounds in clinical use has been heavily investigated ^26, 67–69^. The ever-expanding kinase-substrate interaction map highlights the extensive crosstalk between signaling pathways. Therefore, inhibition of an “off-target” kinase or pathway, cannot be oversimplified and attributed to promiscuity of the SMKI in question; rather it is more likely evidence of an interaction between the intended target and the affected “off-target” signaling node. Two seminal studies collectively examined selectivity of over 170 SMKIs against more than 440 kinases, covering ∼80% of the human kinome. In the first study, Davis et al. compared inhibitor-kinase binding affinities, whereas Anastassiadis et al. used functional kinase inhibition assays in the second study. They determined that while classes of SMKIs may inhibit multiple kinases within a single subfamily, inhibitors are selective against kinases outside that subfamily ^68, 69^. However, a limitation of those studies is that they were carried out using truncated or fused recombinant proteins that may adopt altered conformations in the absence of regulatory domains (i.e., regulatory subunit of PI3K) or binding partners that may influence availability of substrate or ATP binding sites ^26^. Moreover, selectivity of clinically approved SMKIs was validated by super resolution microscopy (dSTORM) to show the superior specificity and selectivity of fluorescently labeled SMKIs like Gefitinib (EGFR inhibitor), over either a fluorescently labeled EGF ligand or an EGFR monoclonal antibody ^70^.

In addition to their therapeutic uses, kinase inhibitors are used as molecular beacons to probe host signaling pathways and delineate how they are regulated by kinases. It is not surprising that NRTKIs which target known effectors of IAV entry have a significant effect on this step of the replication cycle. Indeed, we previously showed that targeting FAK using the pre-clinical inhibitor Y15, led to inhibition of PI3K-mediated endosomal trafficking of virions ^28^. Using the FDA-approved FAK inhibitor DF, we saw comparable effects on actin reorganization and viral entry as we previously observed using Y15 ^28^. This is consistent with the fact that the most prominent effect of DF on viral replication in both A549 and hPCLS was at earlier time-points when reduction in viral entry may have a larger impact than at later time-points. Indeed, we observed a reduction in infectivity in response to DF treatment which is consistent with the observed reduction of viral entry.

Bruton’s tyrosine kinase (BTK) activity regulates survival, proliferation and inflammatory responses in B-cells, as well as epithelial cells ^71^. Cell-specific BTK isoforms have recently been implicated in PI3K and PLC*γ* signaling that are either pro- or anti-apoptotic ^72^. IB and AC are two high-affinity irreversible inhibitors of BTK; IB also inhibits EGFR activity ^73, 74^. IB treatment reduces excessive neutrophil infiltration, acute lung injury (ALI) and subsequent ARDS; ultimately resulting in increased survival of mice severely infected with IAV ^27^. Given that IB inhibits both EGFR and BTK whereas AC selectively inhibits BTK, the IB-specific reduction in viral entry we observed, suggests this effect is mediated largely through inhibition of EGFR signaling. This is consistent with IAV-induced EGFR signaling which facilitates viral entry and activation of downstream pathways (Src, PI3K and ERK) that promote efficient replication ^75^.

BO, along with NI and SA, are second-generation Src inhibitors. BO also inhibits Abl kinase and to a lesser extent BTK. Src orchestrates signaling across multiple pathways that regulate proliferation, survival, cell-cell communication, innate immune responses and apoptosis ^25^. Growth factor RTKs like PDGFR and EGFR induce Src-mediated activation of PI3K/AKT, Ras-Raf-MEK-ERK, FAK, and STAT3 ^76^. Accordingly, Src plays a mostly proviral role during IAV infections that is modulated by the viral NS1 protein ^13, 77^. In contrast, Abl’s role in human IAV infections is not clear; however, Abl inhibition by some avian IAVs results in significant pathology *in vitro* and *in vivo* ^78, 79^. We observed a significant reduction in viral titers following BO treatment of hPCLS and A549 cells which also resulted in a stark reduction in viral entry, largely due to disruption of the actin network. Despite the increase in cell viability during infection, we observed a complete absence of detectible actin filaments suggesting either enhanced depolymerization of F-actin or altered actin dynamics. Consequently, BO inhibition of Src activity can lead to actin depolymerization due to retention of alpha and *β*-catenin at the cell membrane ^80, 81^. To dissect the effect of our NRTKIs on polymerase activity, we employed a polymerase activity reporter system ^82^. Interestingly, in the context of viral infections, we detected higher polymerase reporter activity in NL11 (H3N2)-infected cells. In contrast, significantly higher polymerase activity was detected using NL09 (H1N1) minigenome. Faster kinetics may be more susceptible to NRTKIs as a reduction in replication rate results in exponential differences with time. This suggest that polymerase activity of NL09 is higher than NL11 and that the faster kinetics in virus replication observed in NL11 infected cells may be due to more efficient virus entry, release, or immune evasion than NL09.

In addition to its role in viral entry, we previously demonstrated that FAK regulates the polymerase activity *in vitro* of multiple IAV strains using Y15 as well as dominant-negative kinase mutants ^29^. However, in contrast to our previous studies, we only observed a modest and non-significant effect on polymerase activity following DF treatment. O’Brien et al. showed that Y15 was a significantly more potent and selective inhibitor of FAK activity than DF which also targets the FAK related kinase Pyk2 ^83^. The disparity between a given SMKI’s binding affinity (K_d_) and its functional inhibitory concentrations can also be observed in the case of a single inhibitor targeting multiple kinases. For instance, the K_d_ of the multi-kinase inhibitor Sunitinib for TrkC is 5.1 μM, but a 10-fold lower concentration (0.5 µM) is sufficient to inhibit >97% of its activity. In contrast, Sunitinib’s K_d_ for PAK3 is 16 nM, but not even a 30-fold higher concentration (0.48 μM) has an effect on its activity ^67^. Therefore, the difference in potency of FAK inhibition by DF vs Y15 may account for the limited effect of DF treatment on IAV polymerase activity we observed.

Inhibition of Src by SA, BTK and EGFR by IB, and BTK by AC had less of a significant effect on IAV polymerase activity indicating that the contribution of these kinases to host innate immune responses does not directly affect RNA replication. In contrast, inhibition of Abl and PDGFR*α* by NI treatment had the most significant reduction in IAV polymerase activity that was also strain independent. These data point to a role of PDGFR*α* in facilitating efficient IAV polymerase activity. This is consistent with previous findings that show inhibition of PDGFR*α* by the RTK inhibitor A9, blocks RNA synthesis of all viral RNA species (vRNA, cRNA and mRNA) independently of NFkB signaling ^56^.

The NFkB pathway typically mediates inflammatory/antiviral responses to viral infections and accordingly, it plays a critical role in IAV replication and pathogenesis. Several reports indicate that IAVs modulate antiviral NFkB activity to facilitate viral replication. Inhibition of NFkB results in reduced viral titers partly due to a disruption of vRNP nuclear export ^84–86^. Consistent with previous data, we did not observe a robust induction in NFkB phosphorylation, most likely due to the immuno-suppressive role of the viral NS1 protein ^87^. However, we observed a clear induction of NFkB activation by poly(IC) treatment alone or in combination with IAV infection at 18 h but not 48 h.

In addition to its role in actin reorganization, FAK modulates the cellular immune response by regulating various T-cell-, B-cell-, and macrophage-functions as well as RIG-I-Like antiviral signaling ^88–91^. We previously demonstrated FAK-dependent regulation of NFkB signaling and polymerase activity *in vitro* and NFkB-dependent pro-inflammatory responses *in vivo* ^30^. In that study, FAK inhibition resulted in increased survival, reduced viral load and reduction in a severe infection model. Surprisingly, DF treatment had no significant effect on NFkB phosphorylation; this again, is likely due to the difference in FAK inhibition potency between Y15 and DF. Similarly, none of the other NRTKIs influenced NFkB activation suggesting that the mechanism by which these NRTKIs inhibit virus replication is independent of the NFkB-pathway. Considering the transient and biphasic nature of NFkB activation, we cannot rule out that strain-dependent differences in kinetics did not affect the magnitude or duration of NFkB activation we observed ^92, 93^.

An emerging regulator of IFN and inflammatory responses is STAT3. A wide range of cytokine, growth factor, and RTKs activate STAT3 via JAK1/2/3 and Tyk2-dependent phosphorylation at Y705 (STAT3pY705) ^94^. The role of STAT3 is not fully understood with opposing functions dependent on pathway partner; IL-6 mediated STAT3 activation is proinflammatory while IL-10 mediated STAT3 activation is anti-inflammatory ^94, 95^. Although STAT3 is dispensable for IFN signaling, it is activated by IFN-I and serves as a negative regulator to fine-tune the IFN response (reviewed in ^95^). Because STAT3 activation upregulates anti-apoptotic factors, H5N1 mediated STAT3pY705 allows prolonged viral production through delay of apoptosis; H1N1 is less efficient at STAT3pY705 and triggers apoptosis earlier ^96, 97^. Although the mechanism of differential suppression of STAT3 activation by IAV is not clear, it has been suggested to be mediated by NS1 ^87^. Interestingly, EGFR activation can result in Src/FAK/BTK mediated activation of STAT3, thereby modulating the IFN and proinflammatory responses. As expected of H1N1 and H3N2 infections ^96, 97^, we observed a suppression of STAT3pY705 in untreated cells that was comparable to that observed following treatment with most NRTKIs. Surprisingly, we observed significant suppression of STATpY705 following DF treatment (85-90% of untreated infected cells). Considering that DF inhibits both FAK and Pyk2, it is tempting to speculate that STAT3pY705 requires FAK/Pyk2 activity during IAV infection. Indeed, Pyk2 kinase activity was required to induce EGFR/Src-mediated STAT3pY705 ^98^. Interestingly, STAT3pS727 which is required for full transcriptional activity of STAT3, points to an indirect Pyk2 mechanism possibly through JNK, p38 or ERK activation ^98^.

In summary, we have demonstrated that NRTKs are host cell factors required for efficient IAV replication and represent promising drug targets for the development of the next generation of antivirals. Because these inhibitors target host factors, their therapeutic window is likely to be different than that of virus-targeted antivirals. To our surprise, our tested NRTKIs directly affected steps of the virus replication cycle with limited effects on tested host responses. Importantly, our results were validated using an *ex vivo* lung tissue model from several donors. Given that most of our PCLS were obtained from lung cancer tumor resections, our donors tend to be older, are often smokers and suffer from either COPD or other respiratory pathologies. Although at first glance this may seem like a limitation of our model, we believe that these donors represent the “at risk” populations that would most benefit from IAV antivirals. Therefore, our data obtained from donor PCLS using these already FDA-approved NRTKIs as IAV antivirals is highly applicable to clinical settings. In contrast to virus-directed IAV antivirals which are susceptible to resistance mutations, our data indicate a high genetic barrier for resistance to our tested NRTKIs. This is based on the stability of IAV inhibition after 5 passages under selective pressure by each of our six validated inhibitors. Although we cannot rule out NRTKIs-selected mutations, no resistance/adaptive variants were detected. Additionally, their established safety and bioavailability data further warrants the evaluation of these compounds as potential influenza treatments. Given that IAV infections are typically restricted to the respiratory tract, localized delivery of the kinase inhibitors can further limit potential cytotoxic effects. Finally, the local microenvironment must be considered to elicit a balanced immune response. Likewise, the effect of promising kinase inhibitors on resident and infiltrating immune cells must be investigated to avoid opposite or unintended consequences. Considering that many viruses, including respiratory viruses, utilize the same (or related) host kinases to facilitate their replication and transmission, our studies have broader implications for the potential use of these SMKIs to treat infections by other viruses in addition to IAV infections.

## MATERIALS AND METHODS

### Cells and Viruses

Madin-Darby canine kidney (MDCK) cells were cultured in Dulbecco’s modified Eagle medium (DMEM; Gibco) supplemented with 10% fetal bovine serum (FBS), 100 IU/mL penicillin, 100 µg/mL streptomycin, 2 mM glutamine, and 1% nonessential amino acids (NEAAs). A549 cells were cultured in F-12 K-Nut Nutrient Mix medium (Gibco) supplemented with 10% FBS, 100 IU/ml penicillin, 100 µg/ml streptomycin, and 2 mM Glutamax. All cells were incubated at 37°C and 5% CO_2_.

The pandemic H1N1 strain A/Netherlands/602/09 (NL09) and seasonal strain H3N2 A/Netherlands/241/11 (NL11) influenza viruses were obtained from the Repository of the National Influenza Center at the Erasmus Medical Center in Rotterdam, the Netherlands, and were grown on MDCKs for 48h at 37 °C. Virus stocks and culture supernatants were stored at −80°C until further use. Virus yields were titrated on MDCK cells by 50% tissue culture infectious dose (TCID_50_)/ml method as described by Reed and Muensch ^99^.

### Human precision-cut lung slices (PCLS)

Human PCLS for *ex vivo* studies were generated from lung tissues obtained from patients undergoing surgical operations at Hannover Medical School. Tissues used for PCLS generation that were obtained from lung tumor resections were confirmed as tumor-free by an experienced pathologist. The freshly obtained lung tissues were processed into circular slices that were 300 microns thick and 8 mm in diameter as previously described ^34^. All donors provided informed consent as approved by the Hannover Medical School Ethics Committee (Ethics vote #8867_BO_K_2020). PCLS were maintained in DMEM/F12 medium (ThermoFisher) supplemented with 2 mM of HEPES, 1 × GlutaMAX (Gibco), 100 U/ml penicillin and 100 μg/ml streptomycin in a humidified 37°C and 5% CO_2_ incubator.

### Inhibitors

Small molecule kinase inhibitors (SMKI) were all purchased from Selleckchem (TX, USA). Inhibitors were diluted in DMSO to a stock concentration of 10mM and stored at −20°C upon usage.

### In vitro and ex vivo cytotoxicity assays

*In vitro* cytotoxicity of SMKIs on mock and/or virus infected A549 cells was determined using CellTiter-Glo 2.0 (CTG) Cell Viability Assay (Promega) according to manufacturer protocols.

Cytotoxicity of SMKIs on mock and/or virus-infected PCLS was determined using the LDH-Glo Cytotoxicity Assay (Promega) according to manufacturer’s protocols. Supernatants of SMKI-treated and untreated hPCLS were collected and completely replaced with fresh pre-warmed infection medium containing SMKIs at the indicated concentrations. LDH levels were relative to the positive control (treated with 1% triton-X 100 for 30 min at 37°C).

### Virus infections

A549 cells were plated on the day prior to infection so they were 80-90% confluent on the day of infection. For infections, viruses were diluted in infection medium (F12K containing 0.1% [vol/vol] bovine serum albumin [BSA] and 50 ng/µl TPCK-treated trypsin). The cells were inoculated with the virus at the indicated multiplicity of infection (MOI) for 1h at 37°C. The cells were washed twice with phosphate-buffered saline containing Mg^2+^/Ca^2+^ (PBS+/+) to remove unbound virus and incubated in infection medium at 37°C in the presence or absence of SMKIs at the indicated concentrations. Supernatants were collected at 0, 24, 48, 72 hours post-infection (hpi), and viral titers were determined by TCID_50_ assay in MDCK cells ^99^. GraphPad’s Heatmap (Prism) function was used to visualize the fold-reduction in viral titers.

### Immunofluorescent staining and imaging

To visualize virus infection, infected cells were fixed with 4% paraformaldehyde (4% PFA) for 30 minutes at room temperature (RT), permeabilized with 0.1% Triton X-100 for 15 minutes at RT, washed with PBS and blocked with heat inactivated 5% horse serum in PBS (PBS-HS) at RT for 1h. Cells were then incubated with mouse monoclonal antibodies to IAV nucleoprotein (clone HB65, ATCC) diluted in PBS-HS at 0.2 µg/ml overnight at 4°C under constant agitation. Cells were washed and incubated with AlexaFluor-594 conjugated goat anti-mouse IgG antibody (0.2 µg/ml; ThermoFisher) and NucBlue Live ReadyProbes Reagent (ThermoFisher) for 1h at RT under constant agitation. Cells were washed 3 times with PBS, images were captured using a Leica DMi8 fluorescence microscope and quantitative analysis was performed using ImageJ Threshold, Watershed, and Particle Analyser tools.

Using a counting macro that we adapted from ^100^, we quantified the number of nuclei and the number of separate infected cells by analyzing the RAW image data for each channel (n=4). The nucleus count was used to define the total cell number per 0.6 mm^2^. The NP staining was used to define the number of infected cells per 0.6 mm^2^. The ratio of infected to total cells was used to calculate Relative Infectivity. The total number of cells based on nuclei detected relative to mock-infected cells treated with the respective NRTKI was used to determine Relative Viability. GraphPad’s Heatmap (Prism) function was used for visualization.

### Immuno-histochemistry staining

Mock- and virus-Infected PCLS were inactivated by fixation in 4% PFA/PBS and paraffin-embedded into blocks. Tissue sections (2 µm thick) were cut from the paraffin-embedded blocks and subjected to Hematoxylin & Eosin (HE) staining using standard protocols. Immunostaining for IAV antigen was done using a HRP-conjugated anti-IAV NP antibody. Histological analysis was performed by an experienced pathologist blinded to clinical data and experimental setup using a routine diagnostic light microscope (BX43, Olympus). Representative images were acquired with an Olympus CS50 camera using Olympus CellSens software (Olympus). Semi-quantitative analysis of IAV NP signal was performed for all tested NRTKIs using FIJI image-analysis software.

### Polymerase activity assay

Semi-confluent (∼70–80%) A549 cells (8×10^4^ cells in 24-well plates) were transfected using Lipofectamine LTX with The pPOLI-358-FFLuc reporter plasmid, which encodes a firefly luciferase gene under control of the viral nucleoprotein (NP) promoter (kindly provided by Megan Shaw) ^50–52^; the Lonza pmaxGFP™ expression vector, was used as a transfection control.

For minigenome polymerase activity, a mix of plasmids encoding the PB2, PB1, PA, and NP genes of NL09 or A/NL/213/03 (H3N2) IAVs in quantities of 0.35, 0.35, 0.35, and 0.5 µg, respectively, were co-transfected with the reporter and control plasmids. At 6h post-transfection (hpt), the indicated SMKIs were added at 1x and 0.5x concentrations (see Table 1) and at 30 hpt (24h of treatment), luciferase reporter activity was detected using the One-Glo luciferase assay system (Promega). GFP mean fluorescence intensity (MFI) and luciferase luminescence were measured using a Tecan multi-mode plate reader.

To measure polymerase activity during IAV infection, cells were infected at an MOI of 1 with NL09 or NL11 at 24h post-transfection (hpt) of the pPOLI-358-FFLuc reporter and the GFP plasmids in the presence or absence of SMKIs at the indicated concentrations as described above. At 48 hpt (24 hpi), luciferase reporter activity was detected using the One-Glo luciferase assay system (Promega). GFP mean fluorescence intensity (MFI) and luciferase luminescence were measured using a Tecan multi-mode plate reader.

### Viral entry assay and confocal microscopy

A549 cells were seeded on 12.5-mm coverslips in 24-well plates. On the day of infection, cells were washed 3 times with PBS+/+ and incubated in infection medium in the presence or absence of kinase inhibitors for 2h. The cells were chilled on ice for 15 min and inoculated with virus (MOI=10) in the presence or absence of the indicated SMKI concentrations at 4°C and on ice for 30 min. To limit receptor activation due to continuous viral-receptor engagement/internalization following the 4°C adsorption and to gently warm up the cells, unbound/noninternalized virus was removed by washing the cells twice with RT PBS+/+. The cells were then incubated with prewarmed infection medium containing the respective SMKIs at 37°C for 30 min. Cells were then fixed in 4% PFA for 30 min, permeabilized with 0.1% Triton X-100 at RT for 15 min, washed in PBS, and incubated overnight at 4°C in blocking buffer (PBS-HS). The cells were then incubated with anti-IAV NP antibody (clone HB65, ATCC) diluted in blocking buffer for 1h at RT, washed 3 times with PBS, and incubated for 1h at RT with AlexaFluor488-conjugated donkey anti-mouse IgG secondary antibody (0.2 µg/ml; ThermoFisher) diluted PBS-HS. Cell nuclei and F-Actin were stained with NucBlue Live ReadyProbes (ThermoFisher) and ActinRed-555 ReadyProbes Reagent (ThermoFisher), respectively. Coverslips were mounted with Prolong mounting medium (Invitrogen), and cell images were acquired with a Leica TSC SP5 laser-scanning confocal system mounted on an upright Leica DM6000 CFS using a 63x oil immersion objective. The images were merged and analyzed with Leica LAS software using identical imaging settings across all experiments.

### NRTKI resistance analysis

To assess the resistance barrier for our NRTKIs, we passaged our viruses five times in the presence or absence of submaximal inhibitor concentrations (0.5x; see table 1). The parental viruses were also passaged under the same culture conditions in parallel in the absence of NRTKIs. Semi-confluent MDCK cells (∼10^6^ cells/well in 6-well plates) were infected with the pandemic H1N1 strain A/Netherlands/602/09 (NL09) and seasonal strain H3N2 A/Netherlands/241/11 (NL11) at MOI 0.001. At each passage, the cultures were maintained in 3 ml MDCK infection media at 37°C for 72h, in the presence or absence of the [0.5x]_max_ (see Tab. 1) of respective candidate NRTKIs. Supernatants were harvested, clarified by centrifugation at 500 x g for 5 min at 4°C, and stored at −80°C until titration by TCID_50_ assay on MDCK cells. For the subsequent passage, cells were infected by using virus from the previous passage at MOI=0.001.

### SDS-PAGE and immunoblotting

Proteins were isolated from whole cell lysates using the M-PER Mammalian Protein Extraction Reagent (ThermoScientific). Proteins were quantified by Badford Assay, separated by 8% SDS-PAGE, transferred onto a PVDF membrane and blocked overnight in blocking solution (TBS pH7.6, 0.05% Tween-20, and 5% w/v of nonfat dry milk). Primary antibodies were diluted in blocking solution overnight at 4°C: phos-NFkB p65 (Ser536) (93H1) Rabbit mAb (1:1000) (Cell Signaling), phos-Stat3 (Tyr705) (D3A7) rabbit mAb (1:1000) (Cell Signaling), Influenza A virus NP Antibody (PA5-32242) rabbit pAb (1:20,000) (ThermoScientific). Beta-Actin (BA3R) mAb (1:5000) was used as a loading control. HRP-conjugated secondary anti-rabbit or anti-mouse antibodies (1:20,000) were diluted in blocking solution for 1 hour at room temperature. Proteins were detected by chemiluminescence using the SuperSignal West Pico Plus and SuperSignal™ West Femto Maximum Sensitivity Substrate. Band density was measured using a Li-Cor C-DiGit scanner and analyzed using Image Studio™ (Li-Cor). When necessary, the imaged membrane was subsequently stripped using a mild water-based stripping solution (1.5% glycine; 0.1% SDS; 1% Tween-20; pH 2.2) and restained for total proteins using NFkB p65 (L8F6) mouse mAb (1:1000) (Cell Signaling) or Stat3 (124H6) mouse mAb (1:1000) (Cell Signaling).

### Statistical analyses

Statistical analyses with GraphPad Prism 9 included multiple *t* test, Brown-Forsythe and Welsh’s ANOVA tests and Dunnett’s T3 test for multiple comparisons. Values are represented as means standard deviations (SD) or standard error of the mean (SEM), with a *P* value of 0.05 considered statistically significant (ns = P>0.05; * = P ≤ 0.05; ** = P ≤ 0.01; *** = P ≤ 0.001; **** = P ≤ 0.0001). The performed tests and given significances are provided in the figure legends.

## ACKNOWLEDGMENTS

This work was supported by the Alexander von Humboldt Foundation in the framework of the Alexander von Humboldt Professorship endowed by the German Federal Ministry of Education and Research; and by the European Union’s Horizon 2020 Research and Innovation Program grant number 848166 [ISOLDA]. DJ is supported by a European Consolidator grant, XHale (ref. no. 771883). The author’s would like to thank Regina Engelhardt, Nicole Kröncke, Annette Müller-Brechlin and Christina Petzold-Mügge for their extraordinary technical support.

## AUTHOR CONTRIBUTIONS

HE conceived the project. HE, RM and GR designed the experiments and supervised the project. RM, SS, MB performed the experiments. HE, RM, conducted data analysis. CW, MK and DJ generated and provided human PCLS. CW and MK provided histopathological and immunohistochemical evaluation of PCLS. RM, GR and HE wrote the manuscript with input from all the authors.

## CONFLICT OF INTEREST

The authors declare no conflict of interest.

## Notes

### Competing Interest Statement

The authors have declared no competing interest.

